# Dietary Vitamin B_12_ reduces amyloid-β proteotoxicity by alleviating oxidative stress and mitochondrial dysfunction

**DOI:** 10.1101/2021.02.22.432392

**Authors:** Andy B. Lam, Kirsten Kervin, Jessica E. Tanis

## Abstract

Alzheimer’s disease (AD) is a devastating neurodegenerative disorder with no effective treatment. Diet, as a modifiable risk factor for AD, could potentially be targeted to slow disease onset and progression. However, complexity of the human diet and indirect effects of the microbiome make it challenging to identify protective nutrients. Multiple factors contribute to AD pathogenesis including amyloid beta (Aβ) deposition, mitochondrial dysfunction, and oxidative stress. Here we used *Caenorhabditis elegans* to define the impact of diet on Aβ proteotoxicity. We discovered that dietary vitamin B_12_ alleviated mitochondrial fragmentation, bioenergetic defects, and oxidative stress, delaying Aβ-induced paralysis without affecting Aβ accumulation. Vitamin B_12_ had this protective effect by acting as a cofactor for methionine synthase rather than as an antioxidant. Vitamin supplementation of B_12_ deficient adult Aβ animals was beneficial, demonstrating potential for vitamin B_12_ as a therapy to target pathogenic features of AD triggered by both aging and proteotoxic stress.

## INTRODUCTION

Alzheimer’s Disease (AD), the most common cause of dementia, is a multifactorial neurodegenerative disorder characterized by accumulation of amyloid beta (Aβ) plaques, hyperphosphorylated tau, oxidative stress, mitochondrial dysfunction, and impaired glucose metabolism (Butterfield and Halliwell, 2019; Chakravorty et al., 2019; Lin and Beal, 2006; Long and Holtzman, 2019). Some AD risk factors including genetic predisposition and aging are non-modifiable while other risk factors such as diet can be altered to impact disease onset and progression (Thelen and Brown-Borg, 2020).

Complex diets consist of macronutrients including carbohydrates, fats, and proteins, as well as vitamin and mineral micronutrients. It is challenging to determine which individual nutrients are neuroprotective in humans as well as other mammals due to organismal complexity, genetic diversity, and consumption of complex diets. Indirect dietary effects of gut microbiota, which provide micronutrients to the host, further complicate studies.

The genetically tractable nematode *Caenorhabditis elegans* eats a simple diet of *E. coli* in the laboratory. Consumption of different bacterial strains affects nematode gene expression, metabolic profile, development, fertility, fat storage, and lifespan (Brooks et al., 2009; Cabreiro et al., 2013; Cogliati et al., 2020; Han et al., 2017; MacNeil et al., 2013; Revtovich et al., 2019; Virk et al., 2012, 2016; Watson et al., 2014). With its genetic tools and short lifespan, *C. elegans* is an ideal system for the study of age-related diseases and has been used extensively to identify factors that influence Aβ proteotoxicity (Fang et al., 2019; Han et al., 2017; Hassan et al., 2015; Lublin and Link, 2013; Sorrentino et al., 2017; Teo et al., 2019). Transgenic expression of toxic human Aβ peptides in *C. elegans* body wall muscles generates robust time-dependent paralysis as well as AD-like pathological features including defects in mitochondrial morphology, reduced ATP production, and oxidative stress (Drake et al., 2003; Fonte et al., 2011; Link, 1995; McColl et al., 2012; Sorrentino et al., 2017). Our goal was to use *C. elegans* to define the impact of diet on Aβ-induced proteotoxicity.

We discovered that Aβ-expressing *C. elegans* fed HB101 *E. coli* exhibited delayed paralysis, higher ATP levels, decreased mitochondrial fragmentation, and reduced reactive oxygen species compared to those raised on OP50 *E. coli*. Mild vitamin B_12_ deficiency was observed in animals grown on OP50, but not HB101. We found that B_12_ supplementation delayed Aβ-induced paralysis of animals fed OP50, but did not have an additive impact on those that consumed HB101. The protective effects of vitamin B_12_ required methionine synthase, indicating function as an enzyme cofactor. Vitamin B_12_ supplementation in adulthood was beneficial for B_12_ deficient *C. elegans*, suggesting that administration even late in life has potential as a therapeutic intervention to slow AD progression.

## RESULTS

### Diet alters Aβ-induced paralysis in *C. elegans*

Extraneuronal Aβ plaques (Long and Holtzman, 2019), coupled with genetic evidence (Chartier-Harlin et al., 1991; Goate et al., 1991), suggest that Aβ accumulation is a causal factor in AD development. Transgenic expression of human Aβ^1-42^ in *C. elegans* muscles induces mitochondrial dysfunction, oxidative stress, and robust time-dependent paralysis (Drake et al., 2003; Fonte et al., 2011; McColl et al., 2012). Altered time to paralysis has been used extensively to identify genes and pharmacological agents that influence Aβ proteotoxicity (Cacho-Valadez et al., 2012; Han et al., 2017; Hassan et al., 2015; Lublin and Link, 2013; Sorrentino et al., 2017). While investigating the impact of genetic factors on Aβ-induced paralysis, we noticed that animals fed OP50 B-type *E. coli* consistently paralyzed faster than those given HT115(DE3), an RNAse III deficient K12 derived *E. coli* used for RNA interference experiments. To determine if diet was impacting Aβ proteotoxicity, we gave Aβ animals different *E. coli* diets and discovered that those fed HT115 or HB101, a B x K12 hybrid, exhibited a significant delay in paralysis compared to those that consumed OP50 (Figure 1A and S1A).

**Figure 1:**
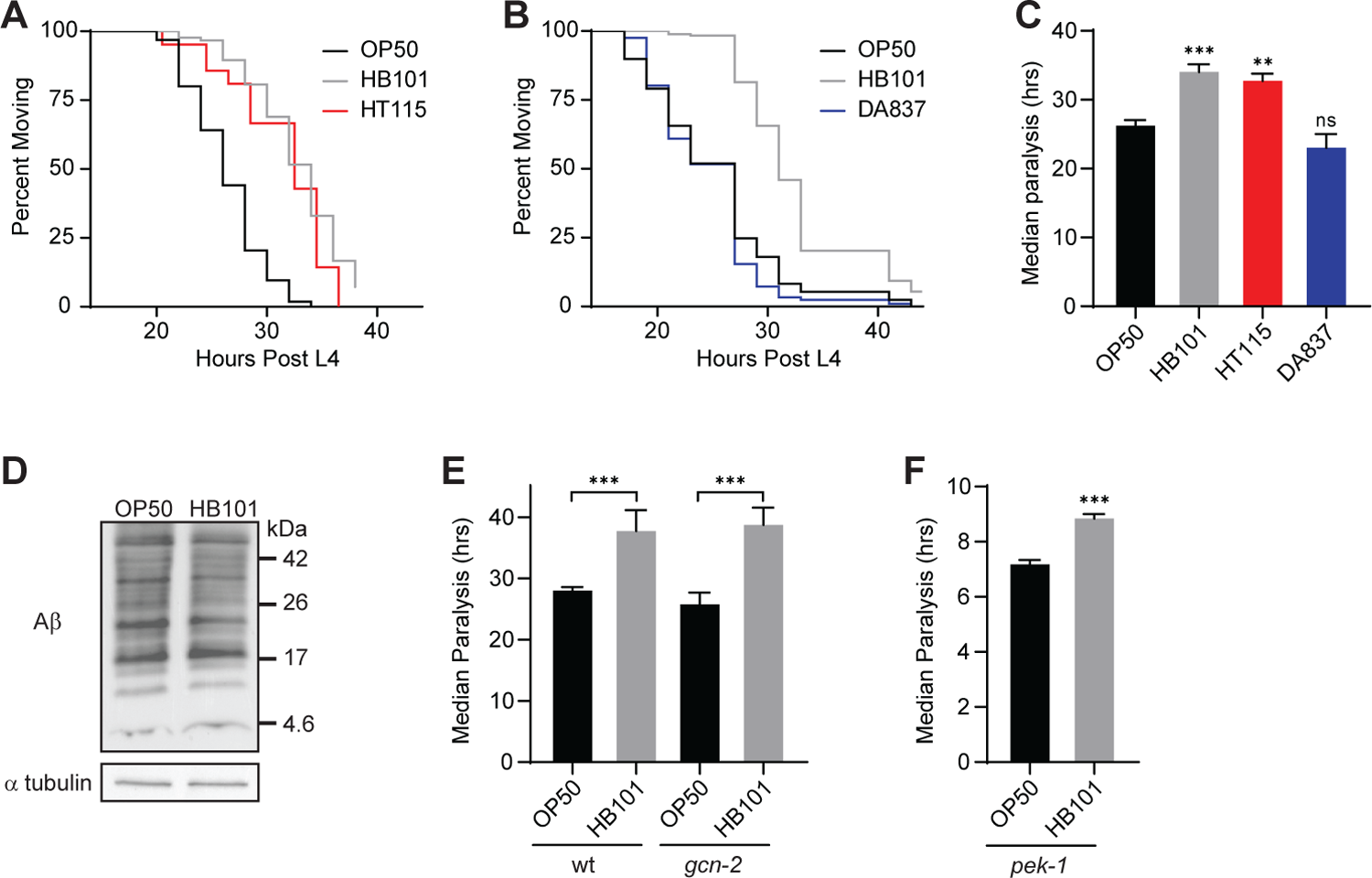
Diet impacts Aβ-induced paralysis without altering Aβ accumulation. (A) Aβ animals fed HB101 (grey) and HT115 (red) exhibited delayed paralysis compared to those on OP50 (black). (B) Aβ animals fed DA837 (blue) and OP50 (black) paralyzed at the same time. (C) Median time to paralysis for Aβ animals fed OP50 (black), HB101 (grey), HT115 (red), and DA837 (blue) *E. coli* (n≥3). (D) Western analysis showed no effect of diet on Aβ accumulation; additional replicates in Figure S1E. (E) Loss of *gcn-2* in Aβ animals raised on either diet did not affect median time to paralysis (n=4). (F) Loss of *pek-1* accelerated Aβ-induced paralysis, but HB101 still caused a delay compared to OP50 (n=3). Error bars show SEM; *p<0.05, **p<0.01, ***p<0.001

Since caloric restriction reduces Aβ toxicity (Steinkraus et al., 2008), we first sought to determine if the altered time to paralysis was due to differences in bacterial growth or *C. elegans* feeding. We found no difference in bacterial concentration on OP50 and HB101 plates (Figure S1B) or pharyngeal pumping rates (Figure S1C) in *C. elegans* grown on the different diets. Aβ animals fed DA837, another *E. coli* B strain that is hard for worms to ingest (Shtonda and Avery, 2006), exhibited paralysis indistinguishable from those given OP50 (Figure 1B,C). Finally, the HB101 diet was still protective in an *eat-2* mutant, which exhibits delayed Aβ-induced paralysis due to dietary restriction (Figure S1D; Steinkraus *et al*. 2008). Together, these results suggest that the diet-induced shift in paralysis was not due to changes in ingestion or dietary restriction.

Reducing Aβ levels delays paralysis (Sorrentino et al., 2017), so we next sought to establish if diet impacted Aβ content. We observed no difference in Aβ accumulation between animals raised on OP50 and HB101 using immunoblotting (Figures 1D and S1E). We also utilized a genetic approach, using two mutants to determine if repression of protein synthesis is required for the HB101 protective effect. The GCN-2 kinase phosphorylates translation initiation factor 2 (eIF2α) during times of nutrient deprivation and mitochondrial stress (Baker *et al*. 2012; Rousakis *et al*. 2013). Loss of *gcn-2* did not affect the dietary shift in Aβ-induced paralysis (Figure 1E). In response to misfolded protein accumulation, the endoplasmic reticulum UPR sensor PEK-1 phosphorylates eIF2α (Shen et al., 2005). *pek-1* mutants expressing Aβ exhibited accelerated paralysis, however, the HB101 diet still caused a delay compared to OP50 (Figure 1F). Our results indicate that the HB101 diet reduces Aβ proteotoxicity without affecting Aβ accumulation.

### Diet impacts mitochondrial morphology and function in Aβ animals

Bioenergetic deficits, which cause synaptic dysfunction in AD individuals, are exacerbated by Aβ accumulation (Chakravorty et al., 2019). *C. elegans* expressing Aβ exhibit decreased ATP levels and defects in electron transport chain complex activity, thus impaired mitochondrial function is a fundamental consequence of Aβ buildup (Fang et al., 2019; Fong et al., 2016; Sorrentino et al., 2017; Teo et al., 2019). Therefore, we tested the impact of diet on ATP levels and found that Aβ animals fed HB101 exhibited a significant increase in ATP compared to those that consumed OP50 (Figure 2A). Diet had no impact on ATP levels in wild-type *C. elegans* (Figure 2A), indicating that the dietary protection was only required for animals under proteotoxic stress.

**Figure 2:**
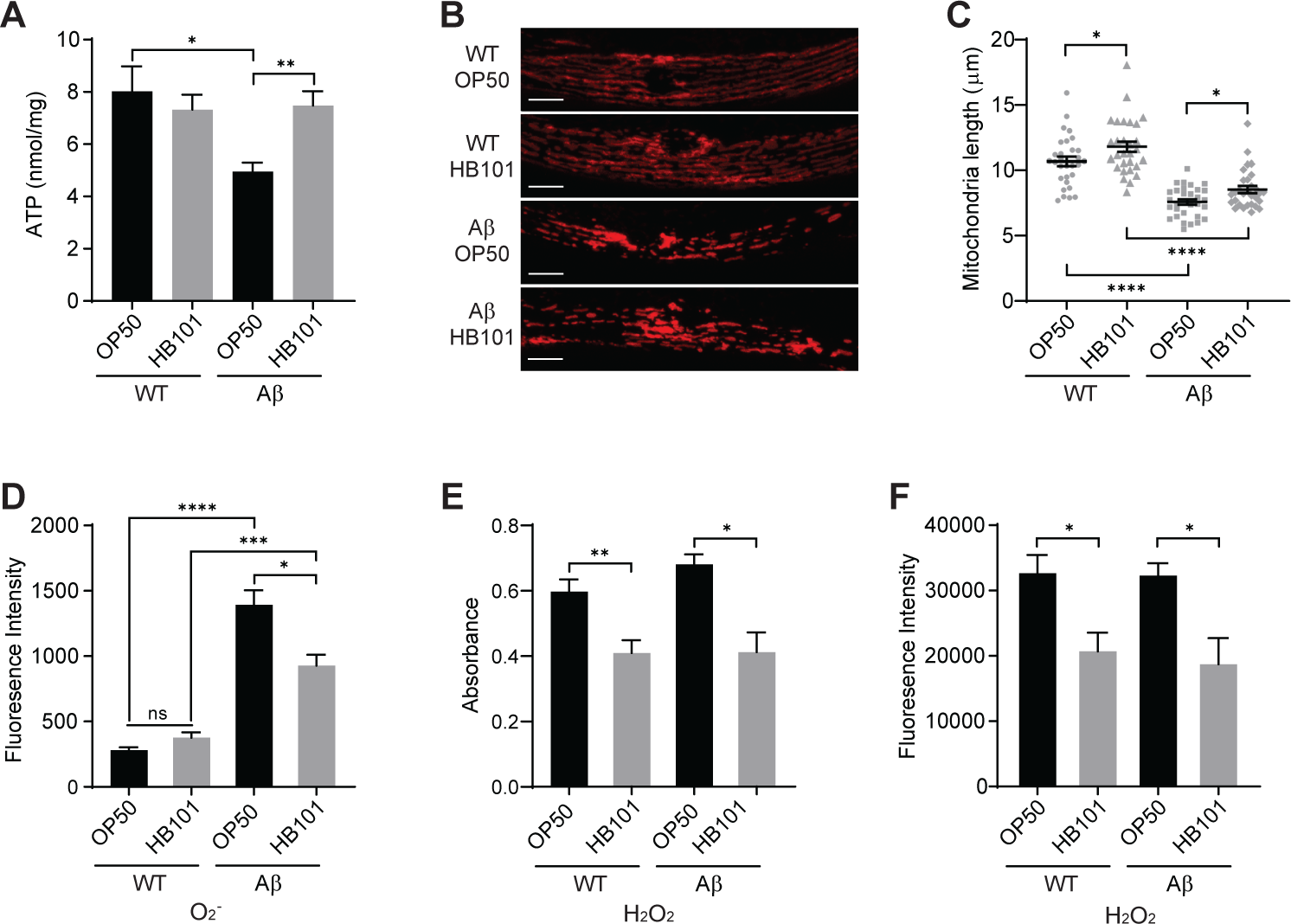
Diet affects mitochondrial function and morphology in Aβ *C. elegans*. (A) ATP levels in Aβ *C. elegans* fed OP50 were reduced compared to wild type (WT); the HB101 diet increased ATP in Aβ animals (n≥8). (B) Representative images of mitochondria, visualized with a TOMM-20::RFP fusion protein, in WT and Aβ animals. Scale = 10 µm. (C) Average mitochondrial length was affected by Aβ accumulation and diet; individual measurements (grey symbols, n≥30) indicated. (D) Superoxide (O^2-^), measured with MitoSox, was higher in Aβ animals fed OP50 versus HB101 (n≥11). (E) Hydrogen peroxide (H_2_O_2_) level, quantified with Amplex Red, was greater in both WT and Aβ nematodes given OP50 (n≥7). (F) H_2_O_2_, measured with H2DCFDA, was increased in WT and Aβ animals fed OP50 (n≥8). Error bars show SEM; *p<0.05, **p<0.01, ***p<0.001

To elucidate the mechanism underlying the diet-induced change in ATP level, we assessed mitochondrial gene transcripts, protein levels, and morphology. Transcript levels of genes required for oxidative phosphorylation (Figure S2A) and mitochondrial protein content (Figure S2B,C) were unchanged between Aβ animals fed the different diets. This suggests that the reduced ATP in animals that consumed OP50 was not due to fewer mitochondria, but rather, diminished function. Aβ expression has been shown to disrupt mitochondrial network integrity, which is important for bioenergetic efficiency (Fonte et al., 2011; Sorrentino et al., 2017). *C. elegans* muscle mitochondria are arranged in a periodic pattern that can be visualized with red fluorescent protein (RFP) tagged TOMM-20 (Wei and Ruvkun, 2020). Mitochondrial fragmentation was observed in Aβ animals compared to wild type (Figure 2B,C). To determine the effect of diet we quantified mitochondria length and found that the HB101 diet reduced fragmentation in both wild type and Aβ animals (Figure 2C). Thus, diet and Aβ expression had an additive effect, indicating that both of these factors impinge on mitochondrial morphology and function.

Oxidative stress, stemming from an imbalance between ROS production and antioxidant defenses, plays a causal role in AD pathogenesis. Mitochondrial fragmentation leads to increased production of reactive oxygen species (ROS) including superoxide (O_2_^-^) and hydrogen peroxide (H_2_O_2_), while Aβ accumulation also promotes ROS formation (Drake et al., 2003; Lin and Beal, 2006; Wang et al., 2014). Therefore, we investigated the impact of Aβ and diet on O_2-_ and H_2_O_2_ levels using ROS indicators. While expression of Aβ resulted in a substantial increase in O_2_^-^, Aβ animals fed HB101 had significantly reduced O_2_^-^ levels compared to those raised on OP50 (Figure 2D).

Both wild type and Aβ *C. elegans* raised on OP50 exhibited significantly higher H_2_O_2_ than those on HB101 (Figure 2E,F). In conclusion, diet and Aβ act in parallel to impact ATP levels, mitochondrial fragmentation, and oxidative stress, three pathological features of AD.

### Vitamin B_12_ protects against Aβ-induced paralysis and mitochondrial dysfunction

OP50 and HB101 differ in carbohydrate content and fatty acid composition (Brooks et al., 2009; Neve et al., 2020; Revtovich et al., 2019). Thus, we considered whether differences in these macronutrients could alter the paralysis of Aβ worms.

Supplementation of plates with glucose (Figure 3A, S3A) or fatty acids (Figure 3B, S3A) did not eliminate the dietary shift in Aβ-induced paralysis, suggesting that these macronutrients are not responsible for the impact of diet. However, Aβ animals raised on OP50 supplemented with glucose paralyzed significantly faster than those fed standard OP50, consistent with reports of excess glucose reducing mitochondrial respiration and shortening lifespan (Lee et al., 2009; Schulz et al., 2007).

**Figure 3:**
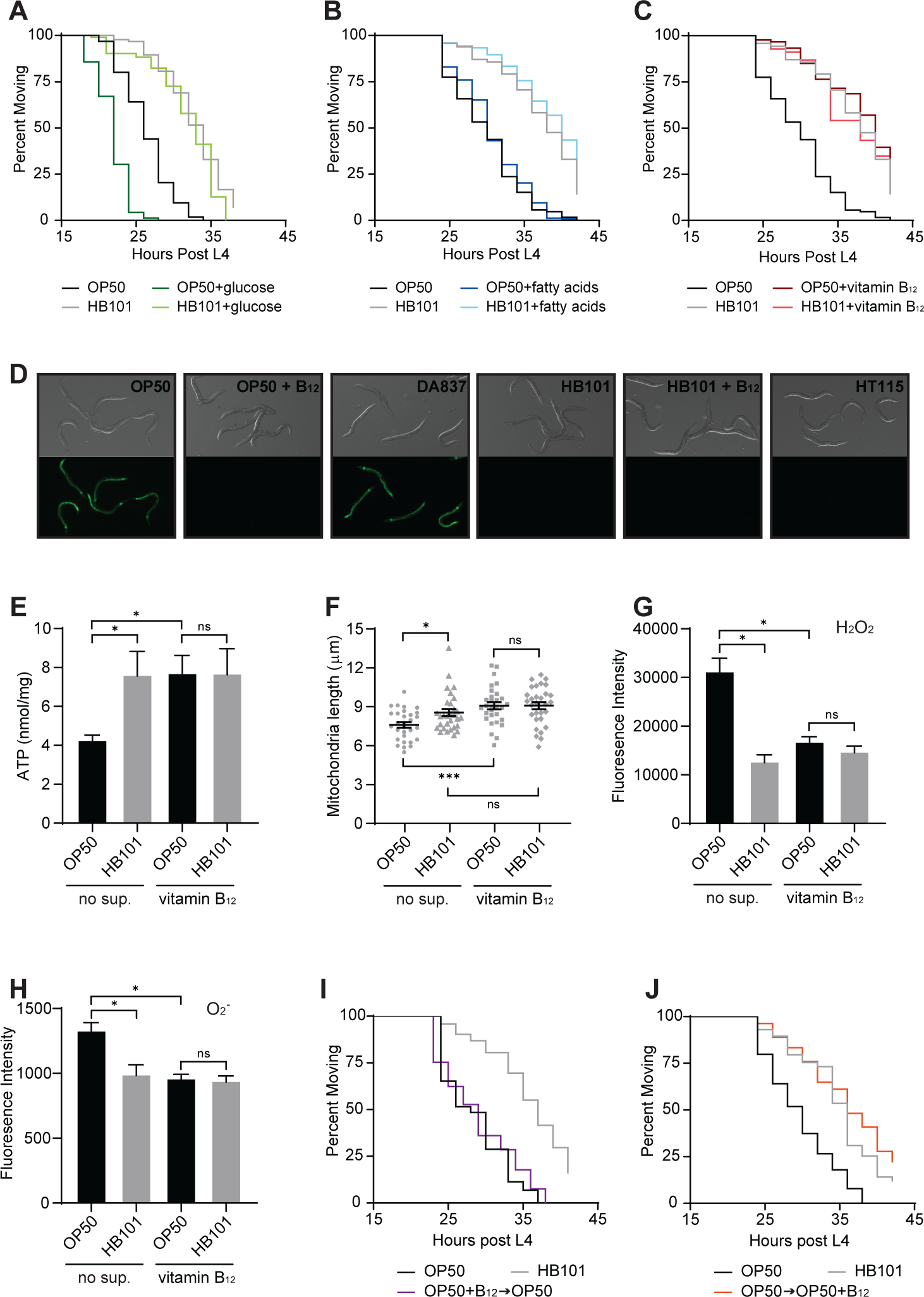
Vitamin B_12_ is the protective factor in the HB101 diet. (A) Supplementation with 10mM glucose accelerated paralysis of Aβ animals fed OP50, but did not eliminate the dietary shift. (B) Supplementation with 0.3mM fatty acids did not affect Aβ-induced paralysis. (C) Supplementation with 148nM vitamin B_12_ eliminated the impact of bacterial diet on paralysis of Aβ animals. (D) *Pacdh-1*::GFP expression was induced in animals fed OP50 and DA837, but not OP50+B_12_, HB101, HB101+B_12_, and HT115. (E) Vitamin B_12_ increased ATP levels in Aβ animals fed OP50 compared to those without supplementation (no sup.; n=6). (F) Vitamin B_12_ increased average mitochondrial length in Aβ animals fed OP50 (n=30). (G) Vitamin B_12_ reduced H_2_O_2_, measured with H2DCFDA, in Aβ animals fed OP50 (n=9). (H) Vitamin B_12_ decreased O^2-^, measured with MitoSox, in Aβ animals fed OP50 (n≥6). (I) Transfer of animals from vitamin B_12_ plates to OP50 at the end of the L4 stage eliminated the protective effect. (J) Transfer of Aβ animals fed OP50 to vitamin B_12_ plates at the end of the L4 stage delayed paralysis. Error bars show SEM; *p<0.05, ***p<0.001

Supplementation of HB101 plates with glucose did not have any effect suggesting that the nutrient in HB101 that protects against Aβ proteotoxicity may also alleviate mitochondrial dysfunction resulting from high glucose. *E. coli* strains also differ in micronutrient content and *C. elegans,* like humans, must obtain several essential vitamins from their diet. Mild vitamin B_12_ deficiency is observed in animals fed OP50, which has reduced expression of the *tonB* transporter that mediates B_12_ uptake (Revtovich et al., 2019; Watson et al., 2014). *C. elegans* vitamin B_12_ status can be assessed using a *Pacdh-1*::GFP reporter, which is expressed in response to propionic acid accumulation resulting from B_12_ deficiency (MacNeil et al., 2013; Revtovich et al., 2019; Watson et al., 2014, 2016; Wei and Ruvkun, 2020). GFP was highly expressed in animals grown on OP50 and DA837, the diets that led to more rapid paralysis of Aβ worms, and B_12_ supplementation suppressed GFP expression (Figure 3D). In contrast, GFP expression was repressed in *C. elegans* that ate the HB101 and HT115 protective diets, suggesting high B_12_ levels (Figure 3D).

Supplementation of OP50 plates with vitamin B_12_ delayed Aβ-induced paralysis, while addition of vitamin B_12_ to HB101 plates did not have any added protective effect (Figure 3C, S3A). This suggests that the dietary shift in paralysis onset was due to differences in vitamin B_12_ availability.

We next sought to determine the impact of vitamin B_12_ on mitochondrial dysfunction, morphology defects, and ROS levels. Aβ *C. elegans* fed OP50 with B_12_ supplementation had significantly higher ATP levels compared to non-supplemented counterparts, while B_12_ had no effect on animals raised on HB101 (Figure 3E). B_12_ supplementation did not affect mitochondrial protein levels (Figure S2B,C). Instead, B_12_ supplementation decreased mitochondrial fragmentation in Aβ animals fed OP50 without impacting mitochondria length in those given HB101 (Figure 3F). Severe B_12_ deficiency in *C. elegans* increases H_2_O_2_ content and decreases antioxidant defense (Bito et al., 2017). We found that addition of B_12_ to OP50 plates significantly reduced H_2_O_2_ and O_2_^-^ in Aβ animals to levels observed in those raised on HB101 (Figure 3G,H). Together these results show that vitamin B_12_ supplementation gives rise to protective outcomes in animals fed the OP50 diet.

### Vitamin B_12_ given during adulthood has protective effects

Subclinical B_12_ deficiency is common primarily in older individuals (Green et al., 2017). Thus, we sought to determine if manipulation of B_12_ availability during adulthood impacted Aβ proteotoxicity in *C. elegans*. Removal of vitamin B_12_ at the beginning of adulthood led to Aβ-induced paralysis indistinguishable from that observed in animals raised on OP50 their entire life (Figure 3I, S3B). This demonstrates that a decrease in vitamin B_12_ levels in adulthood exacerbates Aβ proteotoxicity. Further, we discovered that Aβ animals first grown on OP50 then supplemented with B_12_ only in adulthood exhibited the same delay in paralysis as those fed a vitamin B_12_ rich diet their entire lifespan (Figure 3J, S3B). These results suggest potential for dietary vitamin B_12_ supplementation as an AD treatment even after onset of the disease.

### Protective effects of dietary B_12_ require methionine synthase

Vitamin B_12_ is an essential cofactor for methylmalonyl coenzyme A (CoA) mutase (*C. elegans* MMCM-1), which converts methylmalonyl-CoA to succinyl-CoA in the propionyl-CoA breakdown pathway, and methionine synthase (*C. elegans* METR-1), which converts homocysteine (Hcy) to methionine in the methionine/S-Adenosylmethionine (SAM) cycle (Figure 4A). To define which pathway is required for the B_12_ protective effects, we determined the impact of mutations in propionyl-CoA carboxylase *pcca-1*, which acts upstream of *mmcm-1*, and *metr-1* on Aβ-induced paralysis. Loss of *pcca-1* had no effect on Aβ animals grown on the different bacteria (Figure 4B, S4A), but the dietary shift in paralysis was eliminated in the *metr-1* mutant (Figure 4C, S4B). While OP50 causes mild vitamin B_12_ deficiency, loss of *metr-1* entirely disrupts the methionine/SAM cycle and resulted in an even more severe phenotype (Figure 4C, S4B). Consistent with the behavioral data, loss of *metr-1* in Aβ animals drastically reduced ATP levels and abolished the HB101 protective effect (Figure 4D). H_2_O_2_ and O_2_^-^ content was also the same in *metr-1* Aβ animals raised on OP50 and HB101, though loss of *metr-1* decreased ROS levels, possibly due to reduced oxidative phosphorylation (Figure 4E,F). These results indicate that the methionine/SAM cycle is required for vitamin B_12_ protection.

**Figure 4:**
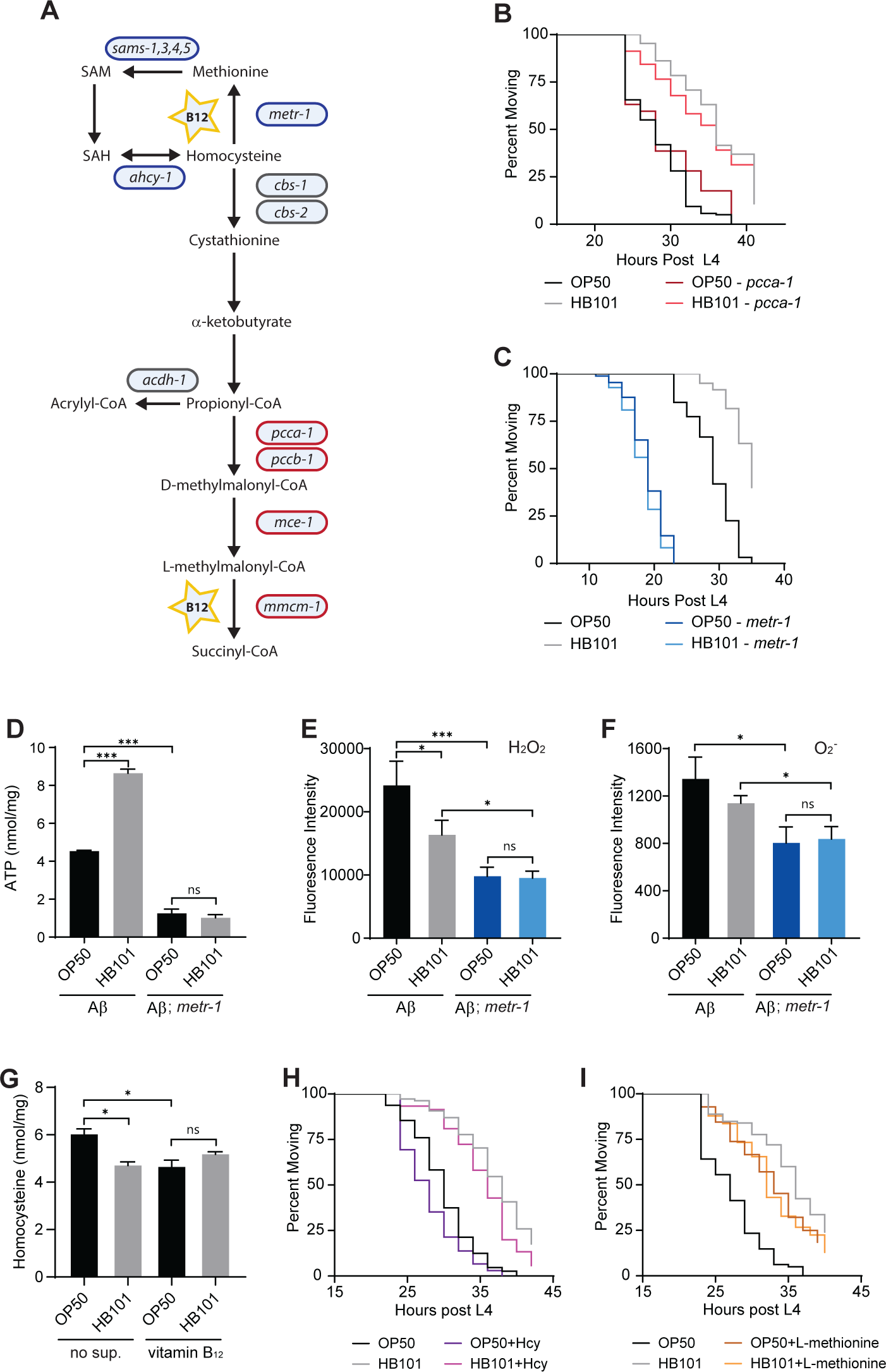
Methionine synthase is required for the protective effects of vitamin B_12_. (A) Diagram of vitamin B_12_ dependent pathways in *C. elegans*. Genes encoding enzymes that function in the methionine/SAM cycle (blue) and the canonical B_12_ pathway (red) are indicated. SAM, S-adenosylmethionine; SAH, s-adenosyl-l-homocysteine. (B) Loss of *pcca-1* did not affect paralysis of Aβ animals. (C) Loss of *metr-1* accelerated Aβ-induced paralysis and eliminated the dietary shift. (D) In *metr-1* mutant Aβ animals, ATP levels were low and unaffected by diet (n=4). (E) H_2_O_2_ levels were unaffected by diet in *metr-1* mutant Aβ animals (n=4). (F) In *metr-1* mutant Aβ animals, O^2-^ levels were unaffected by diet (n=4). (G)Aβ animals grown on OP50 had higher levels of Hcy compared to those raised on HB101 or B_12_ supplemented plates (n=4). (H) Supplementation with 15mM Hcy accelerated paralysis of Aβ animals grown on both OP50 and HB101, but did not eliminate dietary shift. (I) Supplementation with 13.4mM L-methionine eliminated the diet-induced shift in paralysis. Error bars show SEM; *p<0.05, ***p<0.001

B_12_ deficiency reduces methionine synthase activity and can lead to Hcy accumulation. We found that Aβ animals grown on OP50 had significantly increased Hcy, which was reduced by B_12_ supplementation (Figure 4G). Hyperhomocysteinemia is a modifiable risk factor for AD and results in oxidative stress (Bito et al., 2017; Smith and Refsum, 2016). Thus, we sought to determine the effect of Hcy on Aβ-induced paralysis. Hcy supplementation accelerated paralysis of Aβ animals on both diets, but did not impact the dietary shift. This suggests that the increased Hcy in animals fed OP50 did not underlie the effect of low dietary vitamin B_12_ (Figure 4H, S4C). We then tested methionine supplementation and discovered no difference in the time to paralysis between Aβ animals that consumed OP50 and HB101 (Figure 4I, S4C). Methionine supplementation eliminated the dietary shift by having a beneficial effect on Aβ animals fed OP50 and a detrimental impact on those grown on HB101 (Figure 4I, S4C). These data are consistent with a model in which vitamin B_12_ dependent methionine synthase activity controls methionine levels to reduce Aβ proteotoxicity.

## DISCUSSION

Diet is a modifiable risk factor for AD, however, the impact of specific micronutrients on disease onset and progression is difficult to define due to human genetic diversity and diet complexity. Here we used the genetically tractable nematode *C. elegans*, which consumes a simple *E. coli* diet, and discovered that vitamin B_12_ reduces Aβ proteotoxicity. In Aβ-expressing animals, vitamin B_12_ increased energy output, reduced mitochondrial fragmentation, and decreased ROS, but did not impact Aβ levels. There is currently no effective disease-modifying treatment for AD and therapeutics designed to reduce Aβ levels have failed (Long and Holtzman, 2019). Our results suggest that vitamin B_12_ supplementation could be a therapeutic approach to target mitochondrial dysfunction and oxidative stress, pathogenic features of AD triggered by both aging and proteotoxic stress.

Subclinical B_12_ deficiency is common, with a prevalence of 10-15% among individuals over the age of 60 and up to 35% among those older than 80 years (Green et al., 2017) and low B_12_ status may be a modifiable risk factor for AD (Vogiatzoglou et al., 2008). Meta-analyses of clinical trials have suggested that administration of B vitamins does not prevent cognitive decline (Clarke et al., 2014; Ford and Almeida, 2019). However, most participants did not exhibit pre-existing cognitive defects, trial durations were not sufficient to observe decline, and prior B_12_ status was not considered. Our results show that vitamin B_12_ supplementation was beneficial for Aβ-expressing *C. elegans* with mild B_12_ deficiency, but did not offer additional protection for non-deficient animals. Consistent with our work, studies focused on individuals with low dietary vitamin B showed that supplementation with B vitamins preserved cognition (Kang et al., 2008) and slowed brain atrophy (Smith et al., 2010). Thus, the therapeutic potential for vitamin B_12_ is likely to depend on pre-existing B_12_ status. How genetic profile and other components of the complex human diet further impact probability of vitamin B_12_ therapeutic success will need to be resolved.

Dietary B_12_ had no effect on ATP levels in wild type *C. elegans*, suggesting that vitamin B_12_ deficiency is detrimental only during cellular stress. We found that both Aβ accumulation and low B_12_ increased oxidative stress and mitochondrial fragmentation. Since mitochondrial morphology and redox homeostasis are bi-directionally linked, elevated ROS due to Aβ accumulation and mitochondrial fragmentation likely exacerbates mitochondrial defects. Net mitochondrial morphology is defined by fission and fusion as well as removal of damaged mitochondria by mitophagy. Defects in mitophagy decrease ATP levels, elevate ROS, and accelerate Aβ-induced phenotypes (Fang et al., 2019; Palikaras et al., 2015; Sorrentino et al., 2017). Low vitamin B_12_ does not impact mitophagy, but instead promotes mitochondrial fission (Wei and Ruvkun, 2020) and increases oxidative stress (Bito et al., 2017). Thus, vitamin B_12_ and Aβ accumulation may act via different mechanisms to impinge on mitochondrial morphology and thus function.

In *C. elegans*, vitamin B_12_ causes resistance to pathogen stress and this requires methylmalonyl-CoA mutase (Revtovich et al., 2019). In contrast, our work showed that vitamin B_12_ had no effect on *metr-1* mutant Aβ animals, indicating that B_12_ offers protection against proteotoxic stress by acting as a cofactor for methionine synthase.

Methionine supplementation eliminated the dietary shift in Aβ-induced paralysis, but the median time was intermediate to that of animals grown on OP50 and HB101 diets, suggesting that too much methionine can also have a detrimental effect. While methionine restriction causes mitochondrial fragmentation in *C. elegans* (Lin and Wang, 2017; Wei and Ruvkun, 2020), it extends lifespan in other organisms (Grandison et al., 2009; Orentreich et al., 1993). Our work shows the importance of vitamin B_12_ in establishing the proper methionine level to reduce mitochondrial dysfunction and oxidative stress in *C. elegans* under proteotoxic stress.

## SUPPLEMENTAL INFORMATION

Supplemental information includes four figures and can be found with this article online.

## ACKNOWLEDGEMENTS

We thank Jeffrey Caplan from the University of Delaware BioImaging Center for writing the script to measure mitochondrial length. Nematode strains were provided by the *Caenorhabditis* Genetics Center, which is supported by the NIH-ORIP (P40 OD010440). Microscopy access was supported by grants from the NIH-NIGMS (P20 GM103446), NSF (IIA-1301765), and State of Delaware. This work was supported by an NIH-NIGMS INBRE (P20 GM103446) Pilot Project grant and University of Delaware Research Foundation Award #18A00929 (to J.E.T.).

## AUTHOR CONTRIBUTIONS

Conceptualization, A.B.L, K.K., and J.E.T.; Investigation, A.B.L. and K.K.; Writing – Original Draft, A.B.L. and J.E.T.; Writing – Review and Editing, A.B.L., K.K., and J.E.T.; Visualization, A.B.L. and J.E.T.; Supervision, J.E.T.; Funding Acquisition, J.E.T.

## STAR METHODS

### KEY RESOURCES TABLE

See additional file

## CONTACT FOR REAGENT AND RESOURCE SHARING

Further information and reagent requests should be directed to and will be fulfilled by the Lead Contact, J.E. Tanis

## EXPERIMENTAL MODEL AND SUBJECT DETAILS

### Nematode Culture

All *C. elegans* strains were maintained on nematode growth medium (NGM) at 20°C as described (Brenner, 1974). The wild-type strain was Bristol N2; all other strains used in this study can be found in the Key Resources Table. *eat-2(ad465)* was detected with SuperSelective genotyping (Touroutine and Tanis, 2020); standard duplex PCR genotyping was used for all deletion mutants.

## METHOD DETAILS

### Bacterial Strains

*Escherichia coli* strains used in this study include OP50, HB101, HT115, and DA837. Bacterial cultures for assay plates were grown in LB overnight shaking at 37°C to an OD600 of 1.0 (Eppendorf BioPhotometer D30) and seeded onto NGM plates. All plates were dried for three days at room temperature and stored at 4°C before use. Relative amount of bacteria on seeded plates was measured by washing bacteria off plates with M9 buffer following a 24 hour temperature shift to 25°C and OD600 was measured using a BioPhotomer D30 (Eppendorf).

### Aβ-induced Paralysis Assay

Animals were synchronized by a pulse egg lay for paralysis assays, however, we note that the dietary shift in paralysis was also observed with worms synchronized by bleaching or picking of late fourth larval stage (L4) animals to assay plates (not shown). Four adult animals were placed on 6 cm NGM plates seeded with different bacteria for 4 hours and then removed. The animals grew at 20°C and then were shifted to 25°C when the majority reached the late L4 stage, identified by a white crescent with black central dot at the vulva. Starting 12 to 20 hours post temperature upshift, depending on the genotype of the strain being assayed, the number of paralyzed and non-paralyzed animals were counted every two hours. Due to substantial acceleration of Aβ-induced paralysis in *pek-1* mutants, animals were assessed for movement every hour following temperature upshift until all paralyzed. At least 45 animals were assayed per trial and a minimum of three biological replicates were performed per *C. elegans* strain / bacterial condition. Data were used to generate Kaplan-Meier survival plots and determine median time to paralysis (GraphPad Prism).

### Nutrient and Metabolite Supplementation

To make the nutrient supplemented plates for paralysis assays 10mM D(+)-glucose (Fisher Scientific), 0.3mM sodium homogamma linolenate solution (Nu-Chek Prep), 148nM methylcobalamin (Millipore Sigma), 13.4mM L-methionine (Fisher Scientific), or 15mM DL-Homocysteine (Millipore Sigma) were added to autoclaved NGM media at 55°C. These concentrations of glucose (Alcántar-Fernández et al., 2018), fatty acids (Deline et al., 2013), methylcobalamin (Revtovich et al., 2019), L-methionine (Wei and Ruvkun, 2020), and homocysteine (Wei and Ruvkun, 2020) were used in prior work.

Plates were stored at 4°C. Fatty acid supplemented plates were covered with foil to prevent light oxidation. Plates were seeded with the different *E. coli* cultures (OD600 = 1.0) and dried for three days before use.

### Mitochondrial imaging and analysis

Mitochondrial morphology was visualized by imaging *C. elegans* expressing RFP-tagged TOMM-20 (Pmyo-3::tomm-20::mKate2::HA::tbb-2 3’ UTR) 24 hours after temperature upshift of late L4 animals to 25°C. Animals were immobilized with 10 µM levamisole on 3% agar pads and Z-stack images were obtained with a Zeiss LSM880 confocal microscope. Images were analyzed with ImageJ by drawing a ROI around muscles and using the script below to determine mitochondrial length. Values below 1 µm were excluded, then average branch length was calculated. Images from at least thirty animals were analyzed for each condition. The following script was used: run(“Clear Results”); run(“Median…”, “radius=1.5”); run(“Unsharp Mask…”, “radius=2.5 mask=0.90”); setAutoThreshold(“Li dark”); //run(“Threshold…”); setOption(“BlackBackground”, true); run(“Convert to Mask”); run(“Skeletonize (2D/3D)”); run(“Analyze Skeleton (2D/3D)”, “prune=[shortest branch] prune_0 calculate show display”); run(“Summarize”);

### Vitamin B_12_ reporter imaging

Images of P*acdh-1::gfp* expression were taken of adult animals (24 hours post L4) that had been immobilized 3% agarose pads with 10 µM levamisole. All images were collected under identical exposure conditions using a Zeiss AxioZoom V16 microscope with Axiocam 702 mono camera and ZEN 2.3 Digital Imaging System.

### ATP, ROS and Homocysteine Quantification

Approximately 1000 first larval stage (L1) animals synchronized by bleaching were grown on 10 cm plates at 20°C. Once the animals reached late L4, plates were shifted to 25°C for 24 hours (Figures 2 and 3) or 18 hours (Figure 4). Animals were washed 3x with 1x M9 and sonicated on ice with Tris-EDTA buffer (100mM Tris, 4mM EDTA pH 7.75) using a model 150V/T Ultrasonic Homogenizer for 5 minutes, then centrifuged at 14,000 RPM for 15 minutes at 4°C. The supernatant was collected and moved to a fresh tube. ATP quantitation was performed with the ATP Bioluminescence Assay Kit CLS II (Roche Diagnostics) using a Glomax 96 Microplate Luminometer as described (Chaya *et al*, 2021). ATP was normalized to protein content measured with the Pierce BCA protein assay kit (ThermoFisher Scientific). Triplicate technical replicates were performed for each sample; at least four biological samples were assayed for each dietary condition.

For ROS quantification, animals were prepared as for ATP quantification except animals were sonicated on ice in M9 buffer. To normalize samples, the Pierce BCA protein assay kit was used to determine supernatant volume required for 25 µg of protein. Hydrogen peroxide levels were measured using 2′,7′-Dichlorofluorescein diacetate/H2DCFDA (Sigma-Aldrich) as described in (Yoon et al., 2018). Briefly 50 µL of 50 µM H2DCFDA was added to normalized worm samples in black 96 well plates, incubated at room temperature for 6 hours before fluorescence was measured. The Amplex Red Hydrogen Peroxide/Peroxidase Assay Kit (Thermofisher Scientific) was also used to measure hydrogen peroxide levels. Superoxide levels were assessed with the MitoSox Red Mitochondrial Superoxide Indicator Kit (ThermoFisher Scientific) following manufacturers protocols. Fluorescence and absorbance values were measured using the Glomax 96 Microplate Luminometer. Triplicate technical replicates were performed for each sample; at least six biological samples were assayed for each dietary condition.

For homocysteine quantification, animals were prepared as for ROS quantitation. Homocysteine levels in supernatants containing 25 µg of protein were determined using the Homocysteine Assay Fluorometric Kit (Abcam) following manufacturer instructions. Fluorescence was measured with use of the Glomax 96 Microplate Luminometer.

Triplicate technical replicates were performed for each sample; at least four biological samples were assayed for each condition.

### Western Blotting

Eggs from gravid adults were isolated by bleaching and allowed to hatch rocking in M9 buffer overnight. Approximately 5000 starved L1s were pipetted onto each plate the following day. Plates were moved to 25°C when a majority of animals were late L4 and 24 hours later, worms were washed 3x with M9 and the pellet was flash frozen. 100 µL lysis buffer (100mM NaCl, 100mM Tris pH 7.5, 1% NP-40) supplemented with an EDTA-free Protease Inhibitor cocktail tablet (Roche) was added to the pellet, sonicated on ice (Model 150V/T Ultrasonic Homogenizer), and centrifuged 15 minutes at 4°C. The supernatant was transferred to a new tube and 20 µL was set aside for protein quantification (Pierce BCA protein assay kit). 2x protein sample buffer (80 mM Tris-HCl, 2% SDS, 10% glygerol, 0.0006% Bromophenol blue with 10% β-meracaptoethanol and 8M Urea) was added to the supernatant. Samples were heated at 55° C for 5 minutes and 30 µg of protein per sample was run on Tris-Tricine 16.5% precast polyacrylamide gels (BioRad) for Aβ or 10% Tris-Glycine (TGX) precast protein gels (BioRad) for NUO-2. Samples were transferred onto 0.2 µm (Aβ) or 0.45 µm (NUO-2) nitrocellulose membranes (BioRad) for 40 minutes at 70V. Membranes were blocked in 5% non-fat milk in TBS with 0.1% Tween-20 one hour at room temperature. Primary antibody incubation was overnight at 4°C; incubation with a secondary antibody conjugated to horseradish peroxidase was 1 ½ hrs. at room temperature. Chemiluminescence detection used SuperSignal West Pico PLUS (ThermoFisher Scientific) and a Chemidoc MP Imaging System (BioRad). NUO-2 and the α-tubulin loading control were detected together on the same membrane, however, Aβ membranes had to be stripped and reprobed with the loading control antibody to ensure equal protein loading across gels. Antibodies for Aβ (6E10 Biolegend cat#803001, 1:1000), NUO-2 (Abcam cat#ab14711, 1:5000), α-tubulin (Sigma cat#T9026, 1:5000), and goat anti-Mouse IgG-HRP (ThermoFisher Scientific cat#31430, 1:5000) were used in this study. All experiments were done with five biological replicates.

### qRT-PCR

*C. elegans* were synchronized as described for Western blotting. 24 hrs after temperature upshift, animals were washed 3 times with M9 and 1 time with non-DEPC treated RNase free water, then transferred to a microcentrifuge tube. 400 µL of Trizol (ThermoFisher Scientific) was added and samples were flash frozen. RNA was isolated using the RNeasy Mini Kit (Qiagen). Briefly, 200 µL of Trizol were added to thawed samples along with 140 µL of chloroform (ThermoFisher Scientific). Samples were centrifuged, the aqueous layer was removed, and equal volume of 70% ethanol was added. RNeasy spin columns (Quiagen) were used for on-column DNase treatment.

Total RNA was transcribed into cDNA using the iScript cDNA Synthesis kit (BioRad). qRT-PCR was performed with PowerUp SYBR Green Master Mix (Applied Biosystems) using a Quantstudio 6 Flex Real-time PCR System (ThermoFisher Scientific) following the standard cycling mode with an anneal/extend temperature at 58°C followed by a default dissociation step. *act-2* was used as the housekeeping reference. The ΔΔCt method was used to determine relative expression. Triplicate technical replicates were performed for each sample; data presented are from at least three biological replicates per condition.

### Pharyngeal Pumping Rate Measurements

Animals were grown at 20°C to early L4 stage. Using the Zeiss AxioZoom V16 microscope the number of pharyngeal pumps per 30 seconds was counted. At least 28 animals were measured per diet.

### Statistical Analysis

Data were analyzed with one-way ANOVA, performing multiple comparisons with the Dunnett test. Statistical analyses and graphing were performed with GraphPad Prism 7. Significant differences indicated as *p < 0.05, **p < 0.01, ***p < 0.001, ****p < 0.0001.

---

**Supplemental Figure 1:**
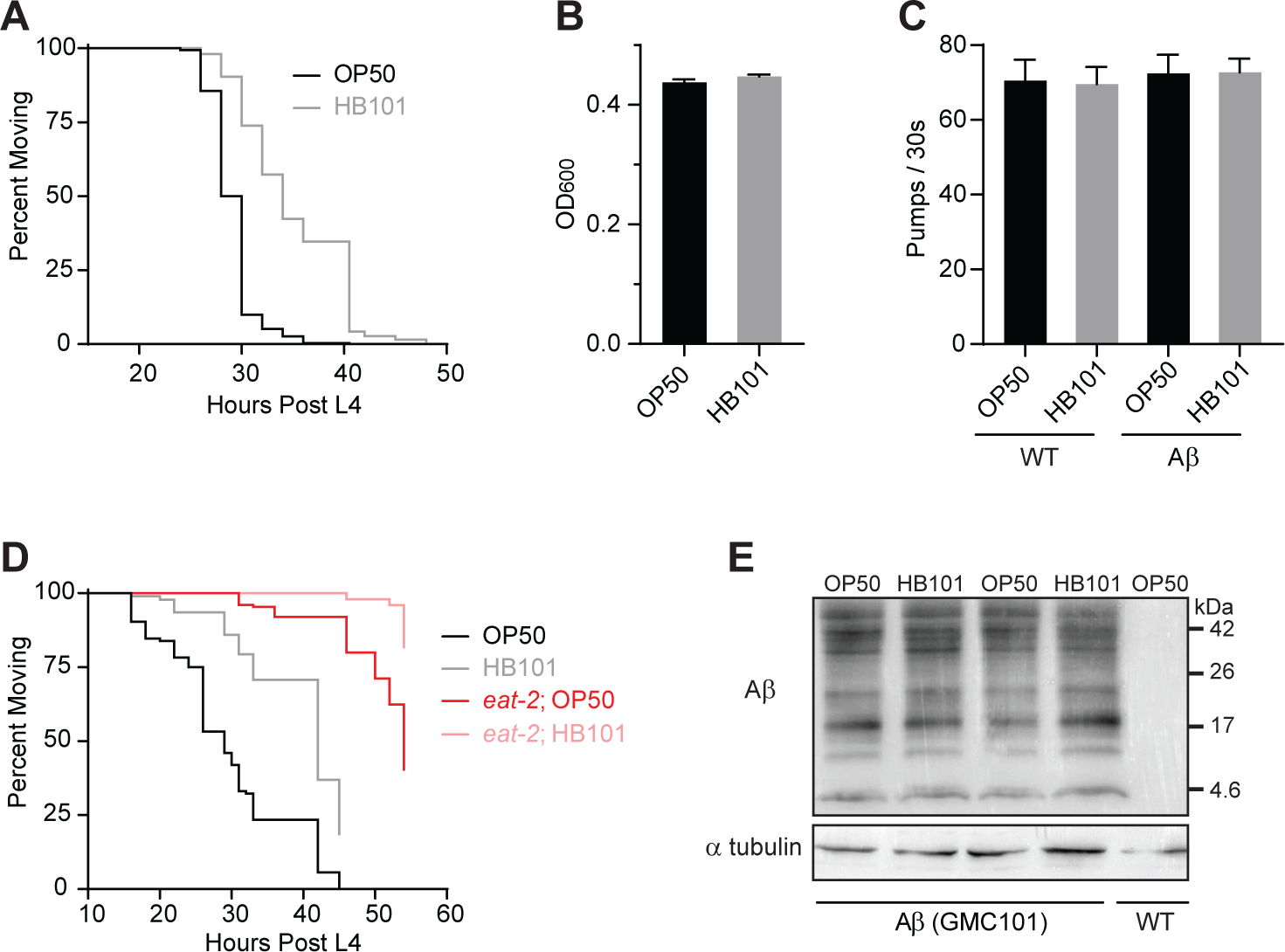
The impact of diet on Aβ-induced paralysis is not due to dietary restriction, variation in ingestion, or differences in Aβ accumulation. (A) Dietary protective effects were not limited to the GMC101 Aβ-expressing strain. CL4176 nematodes, which also express Aβ in the body wall muscles, paralyzed slower when fed HB101 (grey) compared to OP50 (black). (B) OD_600_ measurements showed no significant difference in bacterial growth between OP50 (black) and HB101 (gray) indicating that the Aβ nematodes were exposed to the same amount of bacteria when grown on these diets (n=9). (C) Pharyngeal pumping rates were not different between WT and Aβ nematodes on OP50 (black) and HB101(gray) bacteria (n≥28). (D) Loss of *eat-2* delayed Aβ-induced paralysis in animals grown on both OP50 (red) and HB101 (pink), but HB101 still caused a delay compared to OP50; three biological replicates performed. (E) As in Figure 1D, two additional Western replicates presented here showed no effect of diet on Aβ accumulation; five biological replicates performed in total.

**Supplementary Figure 2:**
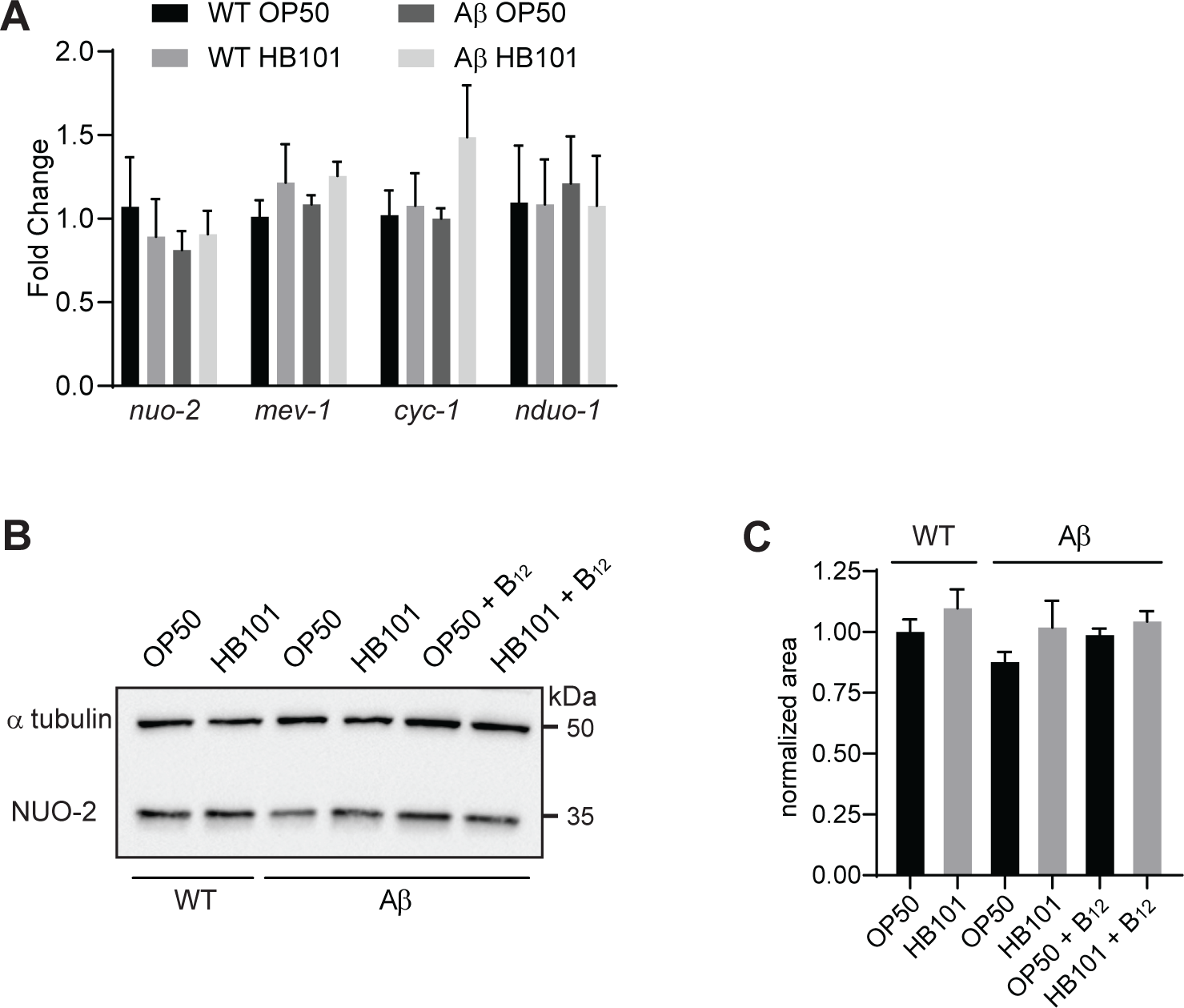
Diet has no impact on oxidative phosphorylation gene transcript levels or mitochondrial protein content. (A) qRT-PCR analysis of transcripts from representative genes required for oxidative phosphorylation (n≥3). No significant differences were observed between wild type and Aβ animals grown on the two different diets. Error bars represent SEM. (B) Western analysis of NUO-2, a component of mitochondrial complex I; the α-tubulin control was detected on the same blot as NUO-2. (C) Quantification of NUO-2 signal, normalized to the a-tubulin control from the same blot. There was no significant difference in NUO-2 protein between animals grown on OP50 and HB101, with or without vitamin B_12_ supplementation. Analysis was performed using Image J on five biological replicates, all run on different gels.

**Supplementary Figure 3:**
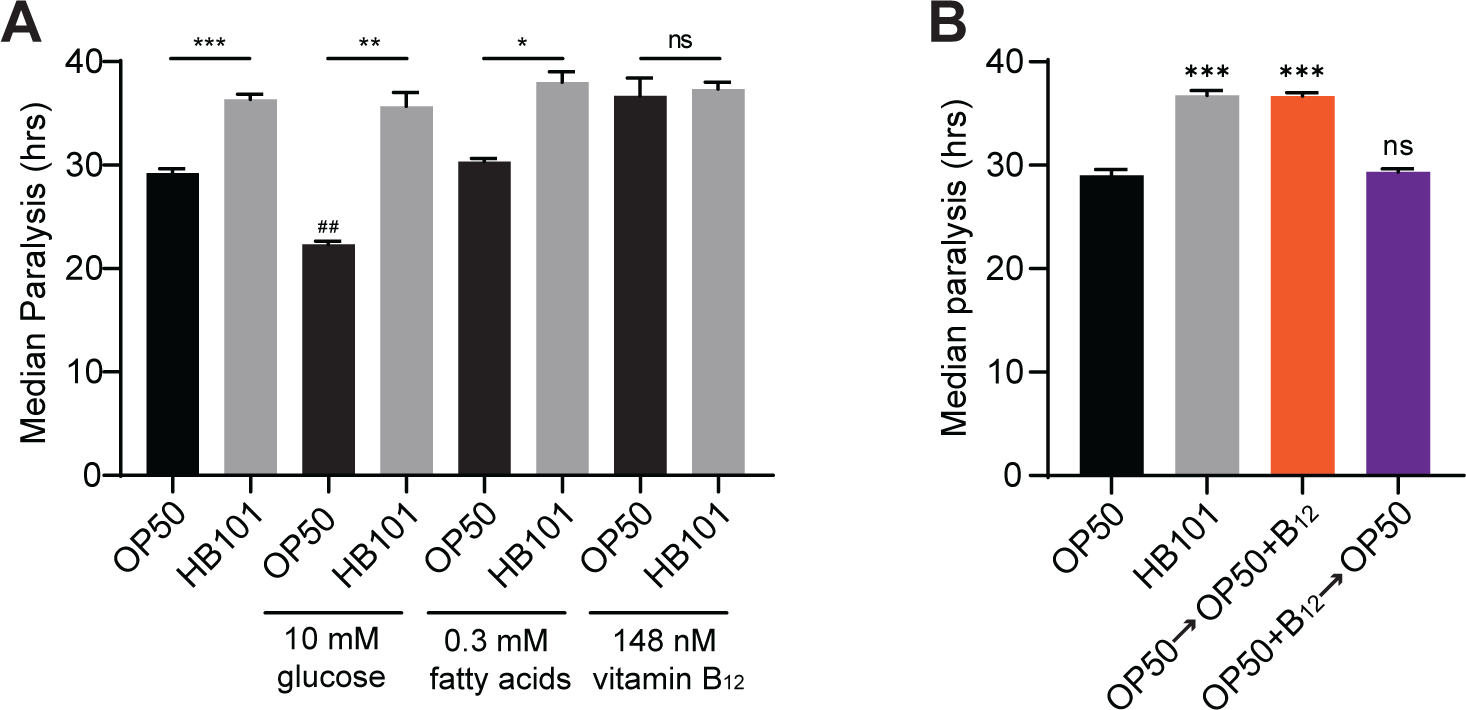
Dietary vitamin B_12_ in adulthood delays Aβ-induced paralysis. (A) Median time to paralysis for Aβ nematodes fed either OP50 (black) or HB101 (grey) on NGM plates with glucose, fatty acids, vitamin B_12_, or no supplementation (n≥3); ^##^p<0.001 compared to OP50 without B_12_. (B) Median time to paralysis for Aβ nematodes fed OP50 without supplementation (black) or with B_12_ introduced (orange) or removed (purple) at the end of the L4 stage (n=3); compare to those raised on HB101 (grey). Error bars show SEM; *p<0.05, **p<0.01, ***p<0.001

**Supplementary Figure 4:**
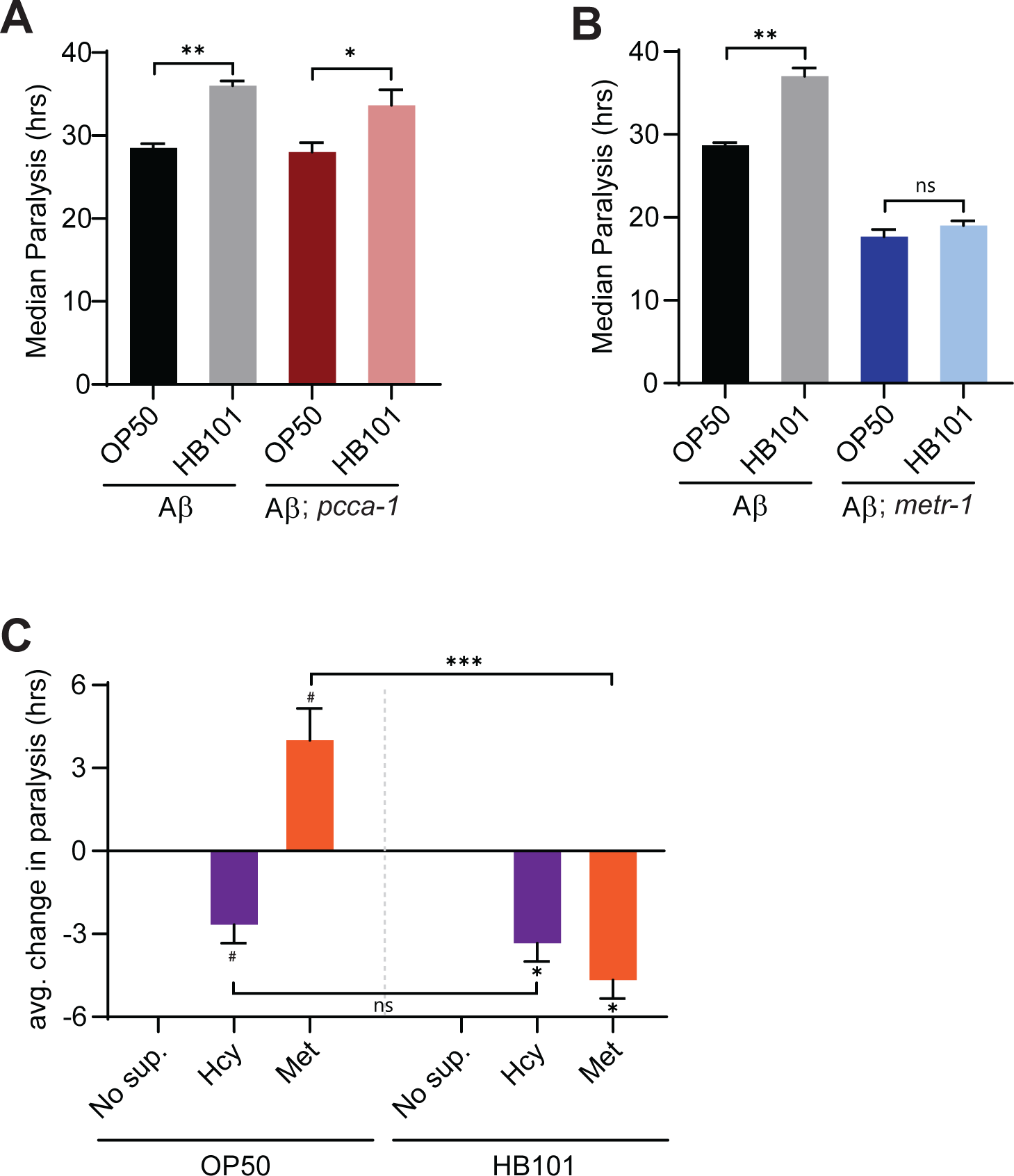
B_12_ deficiency causes defects in the methionine/SAM cycle and an increase in homocysteine levels. (A) Median time to paralysis showed that loss of *pcca-1* had no impact on Aβ-induced paralysis (n=3). (B) Median time to paralysis was not significantly different between Aβ-expressing *metr-1* mutants raised on OP50 or HB101 (n=3). For paralysis assays, error bars show SEM; *p<0.05, **p<0.01. (C) Average change in median time to paralysis (hrs) for Aβ animals on plates supplemented with Hcy (purple) or L-methionine (orange) compared to the respective diet controls (n=3). Error bars show SEM, ^#^p<0.05 compared to OP50 without supplementation, *p<0.05 compared to HB101 without supplementation, ***p<0.001 between animals raised on methionine supplemented OP50 and HB101 plates.

